# Cholinergic Midbrain Afferents Modulate Striatal Circuits and Shape Encoding of Action Control

**DOI:** 10.1101/388223

**Authors:** Daniel Dautan, Icnelia Huerta-Ocampo, Miguel Valencia, Krishnakanth Kondabolu, Todor V. Gerdjikov, Juan Mena-Segovia

## Abstract

Assimilation of novel strategies into a consolidated action repertoire is a crucial function for behavioral adaptation and cognitive flexibility. Acetylcholine in the striatum plays a pivotal role in such adaptation and its release has been causally associated with the activity of cholinergic interneurons. Here we show that the midbrain, a previously unknown source of acetylcholine in the striatum, is a major contributor to cholinergic transmission in the striatal complex. Neurons of the pedunculopontine and laterodorsal tegmental nuclei synapse with striatal cholinergic interneurons and give rise to excitatory responses that, in turn, mediate inhibition of spiny projection neurons. Inhibition of acetylcholine release from midbrain terminals in the striatum impairs action shifting and mimics the effects observed following inhibition of acetylcholine release from striatal cholinergic interneurons. These results suggest the existence of two hierarchically-organized modes of cholinergic transmission in the striatum where cholinergic interneurons are modulated by cholinergic neurons of the midbrain.

## Introduction

The striatum is the main input hub of the basal ganglia. Afferents from the cortex, thalamus and midbrain are widely distributed across its functional domains and together mediate action selection, among other functions. Acetylcholine (ACh) has a powerful influence over striatal circuits. Nicotinic and muscarinic receptors are expressed at pre- and post-synaptic sites in most striatal cell types and their afferents^1–3^, and differentially modulate striatal circuits (see review by^4^). Alteration in cholinergic activity has been shown to have key roles in adaptive behavior. For example, reduced cholinergic transmission impairs the ability to update previous learning and enhances the possibility of interference between novel and old contingencies^5,6^.

Cholinergic markers and released ACh were considered to be exclusively associated with cholinergic interneurons (CINs), which profusely innervate the entire extent of the striatum. While they are more densely concentrated in the matrix of the dorsal striatum^7,8^, their distribution is predominantly random and heterogeneous, thus lacking functional domains^9^. Our recent work demonstrated the existence of an extrinsic source of ACh in the striatum, which originates in the pedunculopontine nucleus (PPN) and the laterodorsal tegmental nucleus (LDT) in the midbrain^10^. PPN innervation of the striatum has been shown to exist in mice, rats and monkeys^11–16^, although its cholinergic nature was only recently revealed. In contrast to CINs innervation, cholinergic innervation arising in the midbrain is topographically organized^10^ and predominantly restricted to the anterior striatum, which receives innervation from prefrontal cortical areas^17^. Thus, the cholinergic midbrain sends topographically organized projections to the entire extent of the anterior striatum, where the rostral segment of the cholinergic brainstem (PPN) preferentially innervates the dorsolateral striatum and the caudal cholinergic brainstem (LDT) preferentially innervates the dorsomedial and ventral striatum. At the synaptic level, PPN and LDT predominantly give rise to asymmetric specializations with dendritic shafts, suggesting excitatory connections, whereas cholinergic interneurons predominantly give rise to symmetric specializations with dendritic spines, suggesting inhibitory connections^10^. The evidence of two sources of ACh in the striatum, each possessing different anatomical characteristics, raises the question of whether they provide differential contributions to striatal circuits.

Cholinergic neurons of the PPN and LDT are phasically activated in response to salient events or changes in brain state^18–20^. Their activation can induce transient fast frequency oscillations in thalamic circuits^21^, which presumably lead to cortical activation and EEG desynchronization. In parallel, cholinergic neurons modulate dopamine mesolimbic circuits that innervate the striatum^22,23^, suggesting that cholinergic neurons have a converging influence on striatal circuits through mesostriatal and thalamostriatal systems^24^. The recent evidence of direct synaptic connectivity with striatal neurons^10^ further suggests that PPN and LDT modulate striatal activity. To understand the impact of the brainstem on striatal function, we used anatomical tracing, *in vivo* electrophysiology combined with optogenetics, and behavior combined with chemogenetics to determine the influence of the cholinergic midbrain on striatal circuits and compared it to that of striatal cholinergic interneurons. Our results reveal two intricately related but distinct modes of cholinergic transmission in the striatum.

## Results

### Midbrain cholinergic neurons contact striatal cholinergic interneurons

PPN/LDT cholinergic neurons preferentially innervate dendritic shafts (76%) in the striatum with a smaller proportion contacting dendritic spines (24%) suggesting a preferential innervation of interneurons over spiny projection neurons (SPNs)^10^. In order to identify the postsynaptic targets of midbrain cholinergic axons, we used a monosynaptic retrograde tracing strategy to label three of the main neuronal populations in the striatum: direct pathway SPNs, indirect pathway SPNs and CINs. Direct and indirect pathway neurons were labeled by injecting a retrograde canine adenovirus (Cav2-Cre) into the substantia nigra pars reticulata (SNR; **Fig. 1A**) or the external globus pallidus (GPE; **Fig. 1B**) of wild-type rats, respectively, thus inducing the retrograde transport and expression of Cre in striatonigral and striatopallidal projection neurons. Subsequently, two floxed viruses were co-injected into the striatum of SNR- and GPE-injected rats to induce the expression of a TVA receptor (AAV-FLEX-TVA-mCherry) and G glycoprotein (AAV-FLEX-G) in direct and indirect pathway neurons. In addition, the same helper viruses were injected in the striatum of ChAT::Cre rats to target CINs (**Fig. 1C**). Two weeks later, a G-deleted pseudotyped rabies virus (SADΔG-eGFP) was injected into the striatum of all three groups in order to infect neurons expressing the TVA receptor *(starter neurons)*. Neurons also expressing the G glycoprotein allowed the transsynaptic transport of the pseudotyped rabies virus, thus labeling those neurons that have monosynaptic connections with Cre-expressing striatal neurons *(input neurons)*^25^. Seven-to-ten days later, the rats were perfused-fixed and their brains analyzed. eGFP-positive neurons were observed in the PPN and LDT of all three groups (**Fig. 1D, F, H; Fig. S1A**), some of which were immunopositive for choline acetyltransferase (ChAT). The total number of ChAT-positive input neurons could not be determined due to interference with the immunohistochemical detection. In some brains one or more of the injections were misplaced and did not show any eGFP-positive neurons thus serving as negative controls. The number of input neurons largely differed between the three experimental groups and CINs-injected rats gave rise to the largest number of PPN labeled neurons (**Fig. 1E, G, I; Fig. S1B**; direct SPNs: 7.33 ± 0.88; indirect SPNs: 12 ± 2; CINs: 24 ± 3.34; Kruskal-Wallis rank-sum test H(2) = 7.482, *P* = 0.0237, *post hoc* two-sample Wilcoxon rank-sum test Z_iSPNs-dSPNs_ = −1.556, *P* = 0.1212, Z_CINs-iSPNs_ = 2.121, *P* = 0.0339, Z_dSPNs-CINs_ = 2.141, *P* = 0.0323). Given the marked differences in the density of SPNs and CINs in the striatum, where SPNs represent about 95% of the total striatal neurons (see review by^26^, we normalized the cell count of input neurons to the number of starter neurons in the striatum. For this purpose, we first analyzed the area of transduction and found that they were not statistically different ([in mm^2^] direct SPNs: 1.04 ± 0.0082; indirect SPNs: 1.36 ±0.01; CINs: 1.62 ± 0.0069; **Fig. S1C-E**; Kruskal-Wallis Rank-sum test H(2) = 2.091, *P* = 0.3515). Then, we counted the number of starter neurons (which correlated with the expected density of each population across similar transduction areas: direct SPNs: 357.55 ± 37.43; indirect SPNs: 367.06 ± 22.57; CINs: 96.32 ±7.68). We then used these numbers to calculate the proportion of input neurons in the PPN based on the number of starter neurons in the striatum for each group (Kruskal-Wallis Rank-sum test H(2)=7.436, *P* = 0.0243). We found that the proportion of PPN input neurons innervating CINs is significantly larger than the proportion innervating either striatonigral or striatopallidal SPNs (**Fig. 1J; Fig. S1B**; *post hoc* two-sample Wilcoxon rank-sum test: Z_dSPNs-CINs_ = −2.121, *P* = 0.0339; Z_iSPNs-CINs_ = −2.141, *P* = 0.0323, Z_iSPNs-dSPNs_ = −1.528, *P* = 0.1212). Similar proportions were observed when WGA-Cre was used instead of Cav2-cre, even though this tracer is expected to diffuse transsynaptically across striatal neurons and therefore overestimate the number of input neurons (**Fig. S1F**; comparison of injections in the SNR, GPE or striatum of WT animals, the latter labeled *all* striatal neurons; Kruskal-Wallis Rank-sum test H(2)=7.395, *P* = 0.0248, *post hoc* two-sample Wilcoxon rank-sum test: Z_dsPNs-all_ = 2.449, *P* = 0.0143; Z_iSPNs-all_ = −2.121, *P* = 0.0339, Z_iSPNs-dSPNs_ = 0.149, *P* = 0.8815). However, the interpretation of the differences between the number of inputs to each striatal cell type is limited to the potential differences in the transduction efficiency of each neuron/pathway, and the results must be taken with caution. For this reason, while it is not possible to estimate the density of innervation of SPNs and CINs from quantifying the number of input neurons in the PPN, our data reveal that a larger number of PPN neurons innervate CINs compared to SPNs, thus suggesting a preferential innervation of PPN neurons to CINs over direct and indirect pathway SPNs.

**Figure 1:**
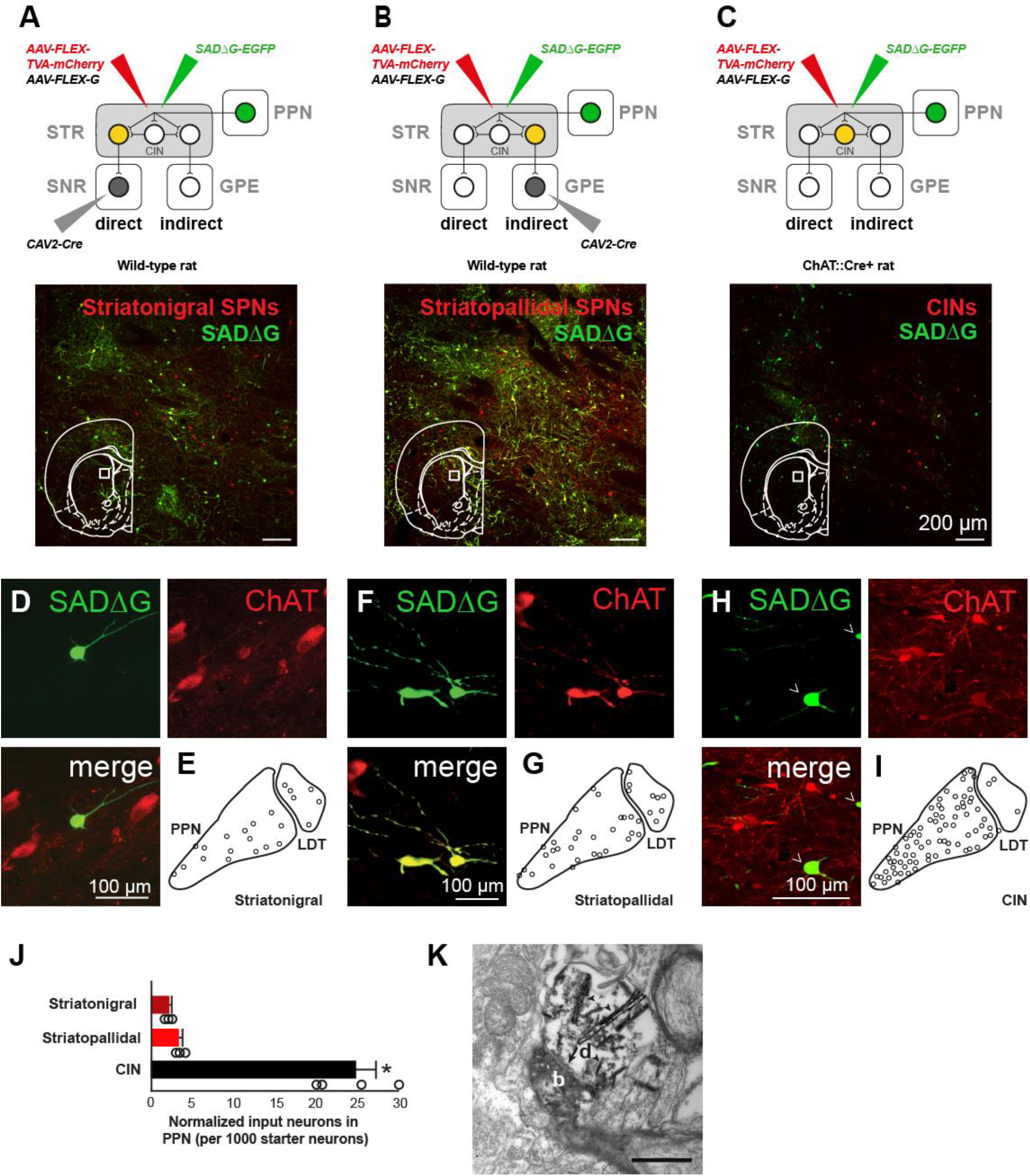
Cholinergic inputs to striatal neurons. (A) Transsynapt¡c labeling of s1r¡aton¡gral SPNs following retrograde Cre transduction in 1he substantia nigra pars reticula1a (SNR) and pseudorabies transduction in the striatum (STR), showing both mCherry-posit¡ve neurons (red, starter neurons) and YFP-positive neurons (green, input neurons). Cholinergic input neurons (D) were observed in the pedunculopontine nucleus (PPN) and (B) Transsynapt¡c labeling of striatopallidal SPN following retrograde Cre transduction in the external globus pallidus (GPE) and pseudorabies transduction in the striatum (STR). Cholinergic inputs neurons (F) were present in the PPN and LDT (G. sum of 3 (C) Transsynaptic labeling of cholinergic intemeurons (CINs) in the ChAT::Cre rat following pseudorabies infection in STR. Cholinergic inputs neurons (H) were present in the PPN and LDT (I, sum of 3 rats). (J) Quantification of inputs neurons in the PPN and LDT (each circle represents one rat, obtained from n = 3 rats in striatonigral and striatopallidal labeling, and n = 4 rats in CINs [an extra animal was added to the analysis]), suggesting that CINs are preferentially (K) Electron microscope image showing an asymmetric synapse in the striatum (black arrow) formed between a cholinergic YFP+· bouton (b; from the PPN) and a CIN dendrite (d). Arrowheads show the accumulation of TMB crystals. Scale bar: K, 500nm. Individual data points and mean *±* SEM are shown. * *P* < 0.05.

Next, to identify synaptic connections between PPN/LDT cholinergic axons and CINs, we used an anterograde strategy based on the transduction of YFP in midbrain cholinergic axons of ChAT::Cre+ rats (n=3) in combination with double immunohistochemistry at the electron microscopic level. PPN YFP-positive axons were converted to a permanent peroxidase reaction using diaminobenzidine (DAB, 0.025%) and nickel ammonium sulfate (0.05%). In addition, CIN cell bodies and processes were immunolabeled with an antibody against ChAT and revealed using tetramethylbenzidine (TMB 0.2%). PPN/LDT axons, identified by the NiDAB reaction product, were observed to make synaptic contacts with dendritic processes of TMB-labeled i.e. ChAT-positive (**Fig. 1K**) and non-labeled structures. Synapses formed with CIN dendrites were identified as asymmetric (Gray’s Type 1) synapses, suggesting an excitatory connection (n = 3), and in line with our previous report identifying the majority of PPN-originated synaptic terminals onto dendritic shafts as asymmetric^10^. These data confirm the transsynaptic retrograde findings and support the evidence of a direct, monosynaptic input from PPN/LDT cholinergic neurons to striatal CINs.

### Differential modulation of striatal neurons by midbrain cholinergic axons

We next tested the effects of stimulating PPN/LDT cholinergic axons on the activity of different types of striatal neurons and compared the effects to the responses elicited by stimulating CINs axons (**Fig. 2, 3**). Cholinergic neurons of the striatum, PPN or LDT were transduced with channelrhodopsin-2 (ChR2) in ChAT::Cre+ rats (AAV2-DIO-EF1a-ChR2-YFP; **Fig. 2A-B**) in order to produce a differential optogenetic activation of midbrain or CINs axons. The spontaneous activity of individual striatal neurons was first recorded *in vivo* in anesthetized animals, then cholinergic axons were stimulated with blue light to activate ChR2 (50ms pulses, 10 Hz) through an optic fiber that was integrated within the recording glass pipette electrode to reduce the spread of the light; the recorded neurons were subsequently labeled with neurobiotin using the juxtacellular method (**Fig. 2C, F, I; Fig. 3A, D, G**;^22^) and their neurochemical nature was confirmed using immunohistochemistry. During urethane-induced slow-wave activity (detected by the electrocorticogram, ECoG), different types of striatal neuron exhibited different firing rates (basal firing rate, SPNs: 1.19 ± 0.13 Hz, n = 91; CINs 2.86 ± 0.37 Hz, n = 53; parvalbumin-expressing interneurons [PV]: 7.24 ± 1.33 Hz, n = 28), in agreement with previous studies^27^. We confirmed that expression of ChR2 in CINs increases their firing discharge during presentation of blue light (**Fig. S2A-D**). The same light stimulation affected neither the firing rate of neurons expressing the reporter alone (AAV-DIO-YFP; **Fig. S2E-G**) nor the activity of their postsynaptic targets (data not shown). Activation of ChR2-expressing cholinergic axons had distinct effects on different subtypes of striatal neurons, and only those recorded/labeled neurons within areas of YFP-transduced axons were observed to respond to the stimulation (**Fig. 2D, G, J; Fig. 3B, E, H**); for this reason, PPN axon stimulation only produced responses in the dorsolateral striatum and LDT axon stimulation only produced responses in the dorsomedial striatum; (**Fig. S3**, see also^10^). In SPNs, all three sets of cholinergic axons produced a significant reduction in the firing rate during the presentation of blue light (**Fig. 2E, H, K**; cluster-based permutation test, 200 permutations, *P* < 0.05). Furthermore, there was no significant effect in the magnitude of the inhibition between PPN, LDT and CINs (% change in firing rates: PPN, −79.58 ± 2.56, n = 29; LDT, −78.39 ± 1.92, n = 19; CINs, −79.24 ± 2.43, n = 43; one-way ANOVA F(2,64) = 0.042, *P* = 0.959). However, the latency of the inhibition was shorter for CINs (0.19 s after laser onset, defined by the inhibition period identified by the permutation test, see blue bar in **Fig. 2K**) compared to either of the midbrain sources, and LDT effects were shorter in duration when compared to PPN or CINs (4.03 s for PPN after laser onset, gray bar, **Fig. 2E**, and 1.92 s for LDT after laser onset, red bar, **Fig. 2H**). Repeated pulses of blue light stimulation in the PPN produced consistent effects across trials and revealed a long-lasting inhibition spanning several seconds (**Fig. S4**). Thus, optogenetic stimulation of cholinergic axons, regardless of the axon origin (i.e. PPN, LDT and CINs), produced a significant decrease in the firing rate of SPNs.

**Figure 2:**
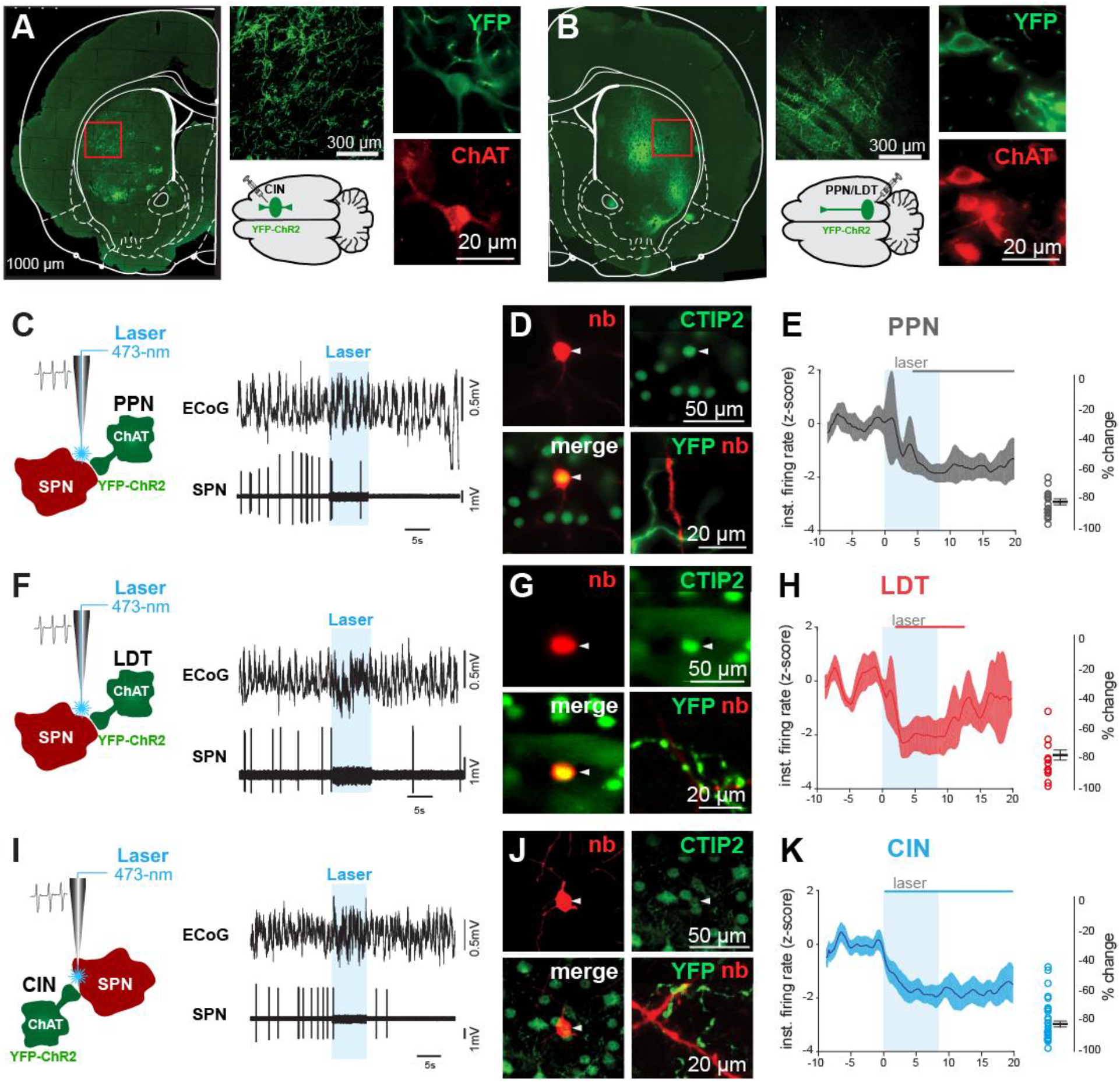
Cholinergic modulation of striatal spiny projection neurons (SPN) (A) Transduction of striatal cholinergic interneurons (CINs) in ChAT::cre+ rats with channelrhodops¡n-2 (ChR2) show dense axonal labeling in the striatum and YFP-posit¡ve somata that were immunopos¡t¡ve for ChAT. (B) Transduction of PPN and LDT cholinergic neurons in ChAT::cre+ rats with ChR2 show patches of dense axonal innervation in the striatum and YFP+/ChAT+ somata in the PPN or LDT. (C) Individual SPN neurons activity was recorded *in vivo* with a glass pipette during optogenetic activation (8s, 10 Hz, 50-ms pulses) of PPN cholinergic axons, and were subsequently labeled with neurobiotin (n = 29 neurons from n = 12 rats). (D) Only neurobiotin labeled SPNs immunopos¡tve for CTIP2 and surrounded by YFP-positive axons were used for further analyses. (E) The normalized instantaneous firing rate of all SPNs that responded to laser stimulation of PPN cholinergic axons shows a slow inhibition during, and after, blue-light stimulation (color line in the top represents the time points during which the responses were significantly different from the baseline; cluster-based permutation test, P < 0.05).(F-H) Same experimental design to assess modulation of striatal SPNs by LDT cholinergic axons (n = 19 neurons from n = 15 rats). LDT cholinergic axon stimulation induced a reduction in the firing rate of SPNs, similar to PPN cholinergic axon stimulation. (I-K) Same experimental design to assess modulation of striatal SPNs by cholinergic axons arising from local CINs in the dorsal striatum (n = 43 neurons from n = 17 rats). CINs axon stimulation induced a reduction in the firing rate of SPNs, similar to the responses of the brainstem.Following cholinergic axon stimulation from each of the three origins, SPNs showed a similar reduction in the firing rate.

**Figure 3:**
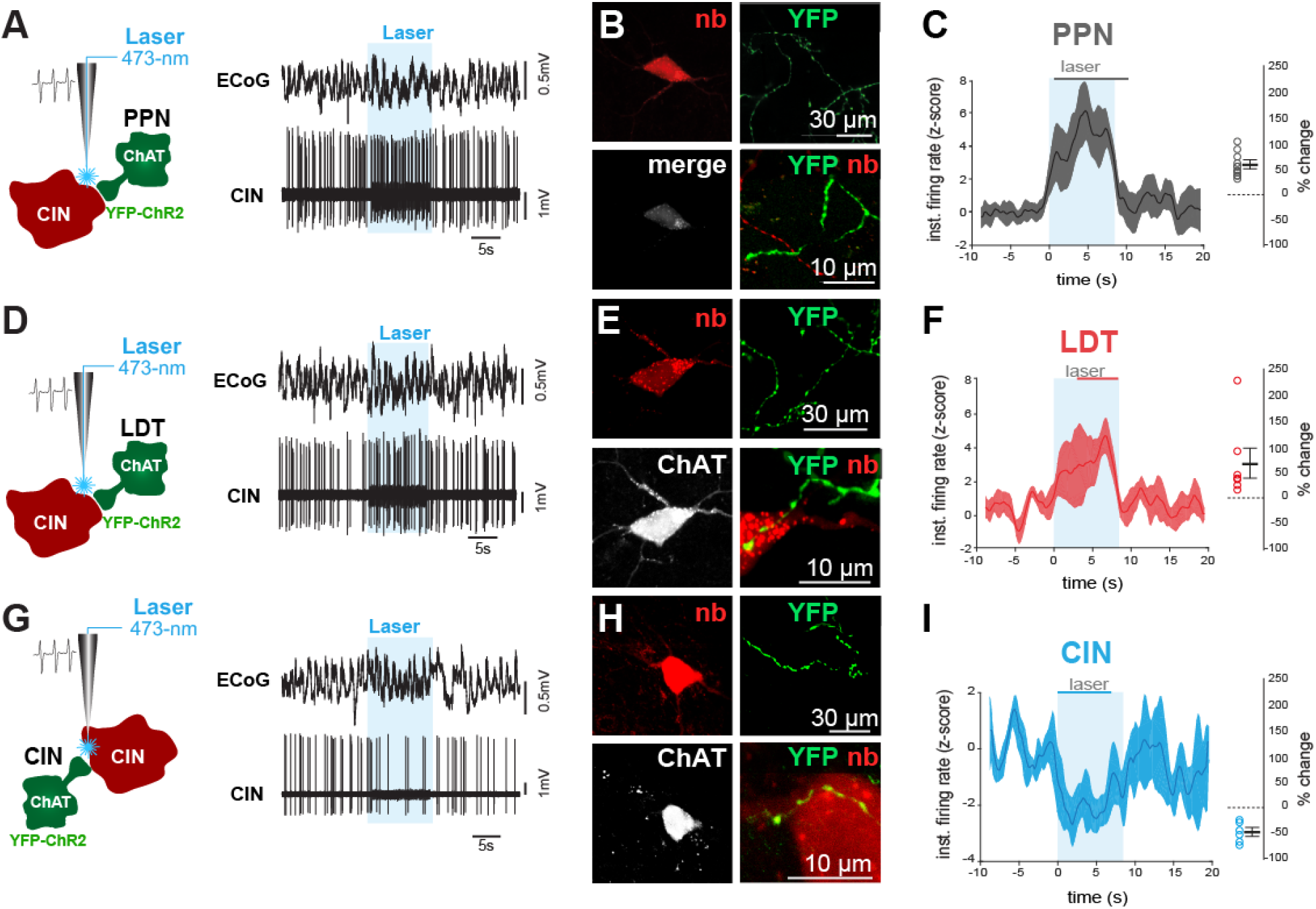
Cholinergic modulation of cholinergic interneurons (CINs) Individual CINs activity was recorded *in vivo* during optogenetic activation (8s, 10 Hz, 50-ms pulses) of cholinergic axons originating in the PPN (A-C; n = 1S neurons), LDT (D-F; n = 13 neurons) or from local CINs (G-l; n = 10 neurons), and were subsequently labeled with neurobiotin. Only neurobiotin-labeled CINs that were immunoposit¡ve for ChAT and surrounded by YFP-positive axons were used for further analyses (B, E, H). The normalized instantaneous firing rate of all CINs that responded to laser stimulation of PPN (C) and LDT (F) show similar increase in firing rate shortly after stimulation, whereas non-transduced CINs (YFP/ChR2-negat¡ve; I) were strongly inhibited during stimulation (color lines in the top represent the time points during which the responses were significantly different from the baseline; cluster-based permutation lest, *P* < 0.05). Similar magnitudes of change were elicited by stimulation of PPN and LDT cholinergic axons.

In contrast to SPNs, the effects on CINs were different depending on the origin of the cholinergic axons: ChR2 stimulation produced excitation of CINs if the axons originated in the midbrain (PPN n = 19, LDT n = 13; **Fig. 3A, C, D, F**) but inhibition if the axons originated in the striatum (**Fig. 3G, I**; cluster-based permutation test, 200 permutations, *P* < 0.05, n = 10; % change in firing rates: PPN, 62.05 ± 7.4; LDT, 52.54 ± 16.2; CINs, −50.17 ± 5.2; one-way ANOVA F(2,41) = 28.19, *P* = 0.00001; Bonferroni *post hoc* analysis: CIN vs LDT *P* = 0.0001, CIN *vs* PPN *P* = 0.0001, LDT *vs* PPN *P* = 1.0). Notably, the latency for producing a significant excitation by PPN afferents was shorter than that provided by LDT afferents (same analysis as above; 0.57 s for PPN after laser onset, gray bar, **Fig. 3C**, and 3.07 s for LDT after laser onset, red bar, **Fig. 3F**). Furthermore, the inhibition of CINs following the stimulation of axons belonging to neighboring CINs was shorter in latency (0.19 s after laser onset, blue bar, **Fig. 3I**) and similar to their inhibitory modulation of SPNs. In contrast to the inhibition in SPNs, which was observed to extend beyond the end of the light stimulation period, the effects on CINs (both excitation and inhibition) were shorter and largely restricted to the stimulation period. Further confirmation of the excitatory effect of the cholinergic midbrain on CINs was observed by analyzing the metabolic activity of CINs (see^28^) following PPN/LDT stimulation. Blue light stimulation of midbrain cholinergic axons in the striatum expressing AAV-DIO-ChR2-YFP increased the immunohistochemical detection of the phosphorylated ribosomal protein S6 in CINs (**Fig. S5**) but not if the axons only expressed the reporter. Our results suggest that midbrain cholinergic neurons are able to activate CINs by increasing their firing rate and increasing their metabolic activity.

Neurons expressing PV did not show a significant effect to the optogenetic stimulation of cholinergic axons originated in either CINs or in the midbrain (n = 28 neurons, % change: PPN = −3.8 ± 64.4, LDT = −9.35 ± 38.4, CIN = 2.01 ± 24.8; one-way ANOVA F(2,26) = 0.16, *P* = 0.8541; **Fig. S6**). While a fraction of PV neurons showed an inhibitory response during the laser stimulation, this response was not consistent across recordings and showed a large variability.

Altogether, the results from the *in vivo* electrophysiology demonstrate that midbrain cholinergic neurons have a differential effect on the dynamics of striatal neurons and their firing rates, inhibiting SPNs and exciting CINs. Furthermore, our data suggest that two functionally distinct sources of ACh operate in the striatum, and that the modulation of CINs seems to be at the center of these differences.

### Striatal circuit effects of midbrain cholinergic activation

Given the evidence of connectivity of PPN/LDT axons with CINs and the differences between PPN/LDT and CINs in their latencies to inhibit SPNs, we then examined whether the inhibitory effects of PPN/LDT on SPNs could be mediated by their excitatory effects on CINs. In a different set of experiments, we transduced PPN/LDT cholinergic neurons with ChR2 and transduced CINs with halorhodopsin (AAV-DIO-mCherry-NpHR3.0; **Fig. 4A**). We recorded the activity of striatal neurons (**Fig. S7**; n = 7 ChAT::cre+ rats) using high-density electrodes (silicon probes) and delivered alternating trains of blue and yellow light, or their combination, in order to activate ChR2 and/or NpHR. Only neurons recorded within the vicinity of midbrain (YFP) and CIN axons (mCherry), as determined by the tracks of the electrode penetration (**Fig. S7A**), were used for further analysis (n = 132 single units). We defined the putative nature of recorded neurons based on their firing rate, action potential duration and coefficient of variation, as previously described (^27,29^; see methods). In line with our results above, putative CINs (pCIN, average firing rate: 1.93 ± 0.38 Hz; n = 8; **Fig. S7C**) were activated by blue light (ChR2; PPN/LTD axons), strongly inhibited by yellow light (NpHR expressed in CINs) and failed to activate during concurrent blue and yellow light stimulation (**Fig. 4B-C, Fig. S7D**; % change of the firing rates during laser stimulation: blue light: 83.10 ± 32.09 % increase; yellow light: 71.19 ± 5.61 % reduction; blue and yellow light: 6.97 ± 22.22 % reduction). In addition, putative SPNs (pSPNs; average firing rate: 1.09 ± 0.09 Hz, n = 33; **Fig. S7B**) were inhibited by blue light (ChR2 expressed in PPN/LTD axons; cluster-based permutation test, 200 permutations, *P* < 0.05; **Fig. 4B-C**). No significant effects on the firing of pSPNs were observed during yellow light stimulation (NpHR expressed in CINs; 11.019 ± 57.61 %; cluster-based permutation test, 200 permutations, *P* = 0.765). During concurrent blue and yellow light stimulation, the inhibitory response of pSPNs was attenuated (cluster-based permutation test, 200 permutations, *P* < 0.05), although it did not disappear (**Fig. 4B-C**; % of firing rates changes during laser stimulation: blue light: −85.07 ± 1.38 %; blue and yellow light: −61.40 ± 15.46 %; paired t-test, t(42) = −9.744, *P* = 2.48×10^-12^) and the duration was markedly reduced (as revealed by the cluster permutation test, compare blue and green bars). These data suggest that the inhibition of SPNs originating from midbrain cholinergic axons is in part mediated by CINs.

**Figure 4:**
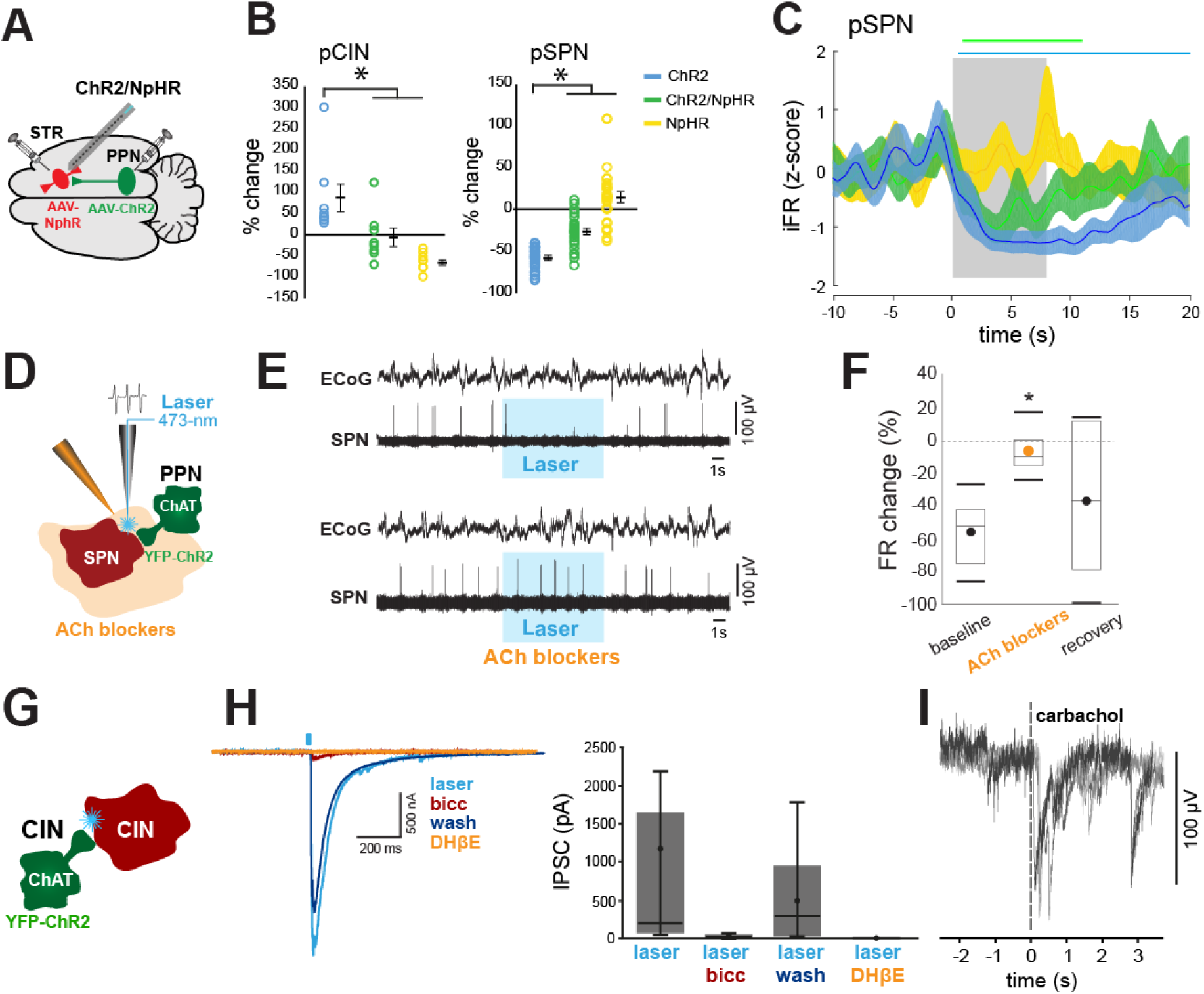
Optogenet¡c and pharmacological dissection of the cholinergic mechanisms that are regulated by the PPN and LDT. (A) Schematic of the *in vivo* optogenet¡c experiment design for extracellular recordings of striatal neurons in ChAT::Cre rats transduced with ChR2 in the brainstem and halorhodopsin (NpHR) in the striatum. (B) Percentage change in the firing rates of pCIN and pSPN during stimulation of brainstem axons transduced wilh ChR2 (blue, 8s, 10 Hz, 50-ms pulses, 5-7 mW), CIN transduced with NpHR (yellow, 8s continuous, 3-4 mW), or both simultaneously (green). The firing rate of CINs was significantly higher during blue light (ChR2 activation} than during yellow (NpHR activation) or blue/yellow light (*P* < 0.Ũ5, see text for statistical values). The firing rate of SPNs was significantly lower during blue light compared to yellow and green light (P < 0.05, see text for statistical values). Individual data points and mean ± SEM are shown. (C) Normalized instantaneous firing rate of all pSPN following stimulation of brainstem axons (blue), inhibition of CINs (yellow), or both simultaneously (green) (color lines in the top represent the time points during which the responses were significantly different from the baseline; cluster-based permutation test, *P* < 0.05). The inhibition of pSPNs by brainstem axons thus seem to depend on the activity of CINs. (D) Schematic of the *in vivo* optogenet¡c and pharmacological experiment design for extracellular recordings of striatal neurons in ChAT::Cre rats transduced with ChR2 in the brainstem and a mixture of acetylcholine antagonists (see text for details) applied through an adjacent glass pipette. (E-F) Individual SPNs that were observed to decrease their firing rate during the ChR2-mediated activation of PPN cholinergic axons, did not decrease their activity in the presence of cholinergic blockers (n = 7 neurons from n = 7 rats; E, representative example; F, group data). (G) Schematic of the *in vitro* optogenelic experiment design for whole cell recordings of CINs in ChAT::Cre mice transduced with ChR2 in the striatum. (H) Whole cell voltage clamp (Vh = −70 mV) recording of identified, non-transduced CINs following optogenetic stimulation of ChR2 (5ms, 450nm) showing consistently IPSCs driven by neighboring CINs axons (n = 9 neurons). IPSCs were strongly reduced by bath application of bicuculline (n = 6 neurons) ora type II nicotinic receptor blocker (DHβE, n = 4 neurons). (I) Representative example of whole cell voltage clamp recording (Vh = −70 mV) of an identified CIN (n = 9 neurons) receiving a local puff of carbachol (100-250 μM, 1 puff per 3 min).

To identify the contribution of ACh to the PPN/LDT-mediated inhibition of SPNs, we used a combined approach using *in vivo* electrophysiology, pharmacology and optogenetics in urethane-anesthetized rats (n = 7), where a small cannula (to deliver acetylcholine receptor antagonists) and an optic fiber (to deliver blue light) were attached to an extracellular tungsten electrode (~200-400 μm from the recording site; **Fig. 4D**). Individual pSPNs (firing rate < 1 Hz and action potential < 2 ms; n = 7 neurons) were recorded during their baseline activity and subsequently during the stimulation of PPN/LDT terminals with blue light to activate ChR2. If pSPNs responded to the stimulation, a cocktail of nicotinic and muscarinic antagonists (100nl in aCSF, 20 mM methyllycaconitine, 40 mM dihydro-β-erythroidine, 40 mM atropine and 100 μM mecamylamine; see^22^) was infused and the response to the laser was tested again 15 min after the infusion (**Fig. 4E**). pSPNs that decreased their firing rate as a result of the blue light stimulation (−56.25 ± 7.47%) showed a diminished inhibition in the presence of cholinergic blockers (−7.2 ± 4.9%; 15 min after drug delivery; **Fig. 4F**). The inhibition to the laser was partially recovered ~45 minutes after the drug application (−36.15 ± 16.37%; one-way ANOVA F(2,20) = 5.24, *P* = 0.0161; Bonferroni *post hoc* analyses: before *vs* during *P* = 0.014, before *vs* after *P* = 0.611, during *vs* after *P* = 0.221).

Finally, to determine the effects of the optogenetic activation of cholinergic axons, we used an *ex vivo* approach. Cholinergic neurons of the PPN or striatum of ChAT::Cre+ mice were transduced with ChR2-YFP and recorded *in vitro* (**Fig. 4G**). We observed an inhibitory response in YFP-negative CINs when axons of neighboring YFP-positive CINs were activated (blue laser, 5 ms pulse; **Fig. 4H**), in line with our *in vivo* experiments (see **Fig. 3**) and with previous reports 30. This inhibitory response was abolished in the presence of bicuculine or DHβE (**Fig. 4H**), suggesting a disynaptic mechanism mediated by GABAergic interneurons^31^. We were unable to detect any effect of PPN/LDT cholinergic stimulation in the slice (as also observed in other PPN targets, such as in the thalamus [unpublished data], the VTA [^22^], or even locally in the PPN; see also^32^), probably due to a low preservation of PPN cholinergic axons in the slice. Nevertheless, local administration of carbachol to CINs in the presence of glutamate blockers (CNQX and AP5 10μM) and a muscarinic blocker (atropine 0.5μM) produced large excitatory currents in 4 of the 11 CINs recorded, possibly mediated by nicotinic receptors (**Fig. 4I**; see also^33^). Our results altogether suggest that PPN/LDT cholinergic axons inhibit SPNs through a combined effect that is partly mediated by CINs, and directly excite CINs through a potential nicotinic effect.

Additional mechanisms are likely to contribute to these circuit effects, such as the pre-synaptic activation of corticostriatal or thalamostriatal terminals^34,35^, or the activation of other types of GABAergic interneurons^36^. Further experiments are necessary to understand the full extent of the midbrain effects on striatal circuits.

### Encoding of behavior by cholinergic systems in the striatum

Cholinergic transmission in the striatum has been associated with updating of action-outcome associations. CINs have been shown to facilitate the integration of new learning into old strategies, whereas cholinergic PPN neurons seem to be involved in behavioral shifting and updating the behavioral state triggered by changing contingencies^24^, thus having a seemingly convergent function. In order to interrogate the contribution of the midbrain cholinergic system in striatal-dependent behavior, we used a chemogenetic strategy to inhibit the local release of acetylcholine in the striatum^37^ during the acquisition of an instrumental lever-press task that reveals action-shifting between goal-directed and habitual strategies^38–41^. Thus, ChAT::Cre+ and wild-type rats were injected with AAV-DIO-hM4Di-HA-mCherry into the PPN, LDT, dorsolateral or dorsomedial striatum (**Fig. S8A**). Bilateral cannulas were implanted in the dorsolateral (for DLS and PPN groups) or dorsomedial striatum (for DMS and LDT groups) for intracerebral delivery of clozapine-N-oxide (CNO; **Fig. S8D**), which binds and activates the transduced hM4Di receptors (associated with the Gi protein) and significantly reduces cell firing in cholinergic neurons, as demonstrated in slice recordings (paired t-test, t(6) = 3.677, *P* = 0.0104, **Fig. S9**). Before each training session, rats received intrastriatal infusions of CNO (1.5 μM, 250 nl, 30 min before), which was calculated to diffuse 300-500 μm from the tip of the cannula, as revealed by fluorogold injections at the end of the experimental procedure (**Fig. S8 D-E**). Rats were trained to press a lever to obtain a reward in a random ratio (RR) schedule and then switched to a random interval (RI) schedule (**Fig. S10A**); the former has been associated with the formation of goal-directed behavior whereas the latter has been associated with the formation of habitual behavior (**Fig. S10B-C**).

The control group consisted of wild-type rats receiving the same manipulations (i.e., viral injection, cannulation, and CNO delivery) and training as the experimental group. Animals showing histological signs of striatal lesions in any group were not considered for further analysis. No differences due to CNO (versus saline) infusion were observed in any group [WT and ChAT::cre+ rats, each virally transduced in the DLS, DMS, PPN or LDT] in locomotor activity (**Fig S10D-E**), evaluated as total distance travelled (two-way ANOVA group x drug: F_group_ (3,35) = 1.69, *P* = 0.1924; F_drugs_(1,35) = 1.86, *P* = 0.1840; F_interaction_(3,35) = 0.92, *P* = 0.4461) and distance in center of the open field (two-way ANOVA group x drugs condition: F_group_ (3,35) = 1.06, *P* = 0.3820; F_drugs_(1,35) = 0.61, *P* = 0.4429; F_interaction_(3,35) = 2.48, *P* = 0.0815). No changes were detected either in sugar consumption (two-way ANOVA groups x drugs: F_group_(4,99) = 1.94, *P* = 0.11, F_drugs_(1,99) = 0.001, *P* = 0.96, F_interaction_(4,99) = 0.08, *P* = 0.99), suggesting that midbrain cholinergic terminals targeting other structures were not affected (see^42^). During training, the number of lever presses during RR showed incremental changes in all groups (**Fig. 5A**), whereas during RI they remained constant (**Fig. 6A**), consistent with previously reported data^43,44^. All animals were then tested in an outcome devaluation task, consisting of two counterbalanced sessions carried out over two consecutive days: a ‘valued’ session where rats were fed rat chow but no sugar pellets (the instrumental outcome) immediately before testing (maintaining a high motivational state for the outcome) and a ‘devalued’ session where rats were fed sugar pellets before testing (thus devaluing the instrumental outcome). While goal-directed behavior is expected to be sensitive to the motivational changes of the devalued session, resulting in a reduction in the number of lever presses, habitual behavior is not expected to be affected by pre-exposure to the instrumental outcome and therefore lever pressing in the devalued session should not be significantly reduced^38,39,45^.

**Figure 5:**
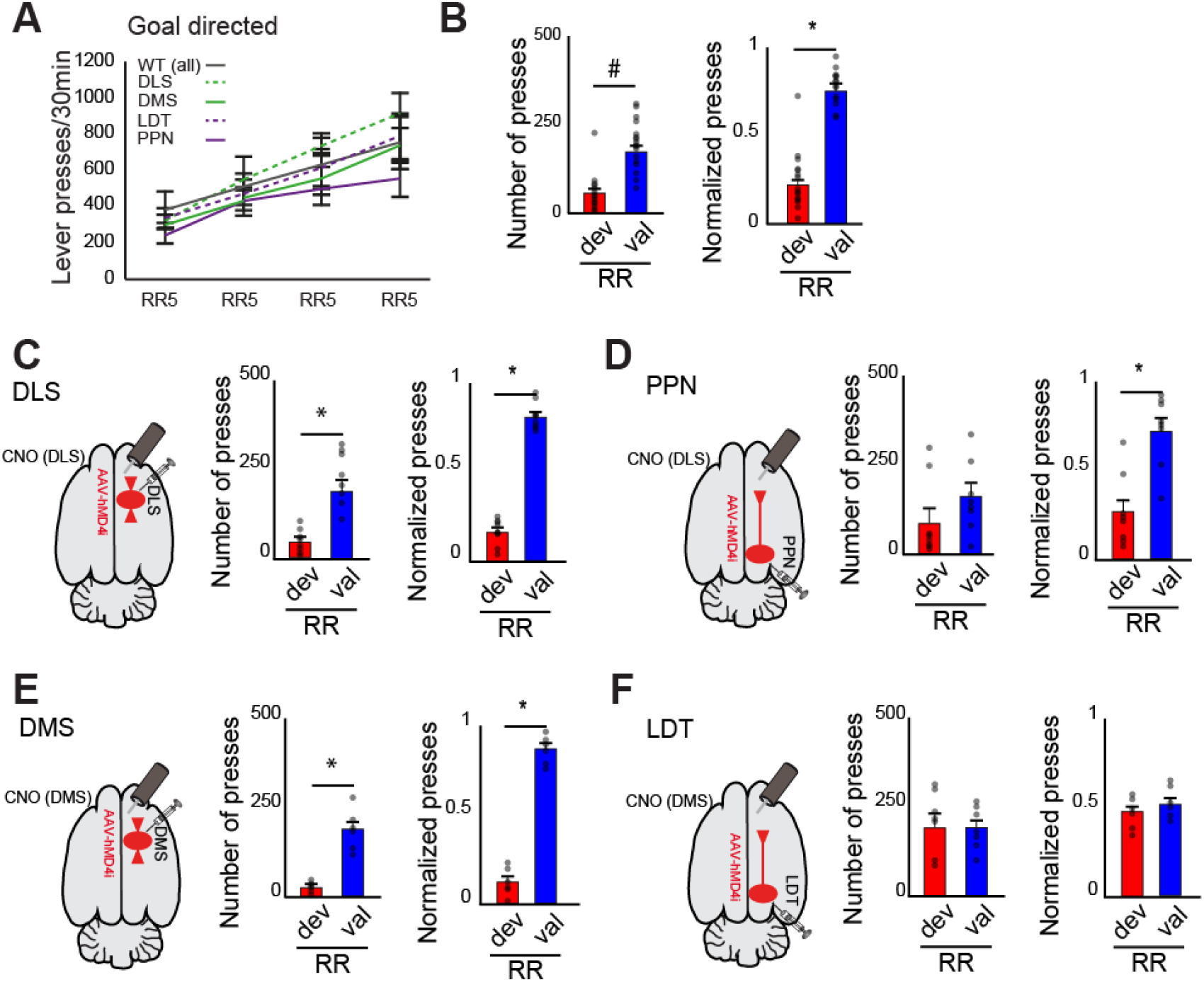
Blocking of LDT cholinergic transmission in the striatum impairs goal-directed action control. (A) Lever pressing during acquisition of goal-directed behavior shows no significant difference between groups (see text for details). (B-F) Number of presses and normalized lever presses during outcome devaluation testing across valued (val) or devalued (dev) states in random ratio schedule (RR; goal-directed). Control animals (WT, B) and ChAT::Cre animals were injected in the DLS (C), PPN (D), DMS (E) or LDT (F). During RR, significant differences in goal-directed devaluation were observed in all groups except LDT, suggesting that inhibition of LDT axons prevents animals from switching to goal-directed behavior. Individual data points and mean ± SEM are shown. * *P* < 0.05.

**Figure 6:**
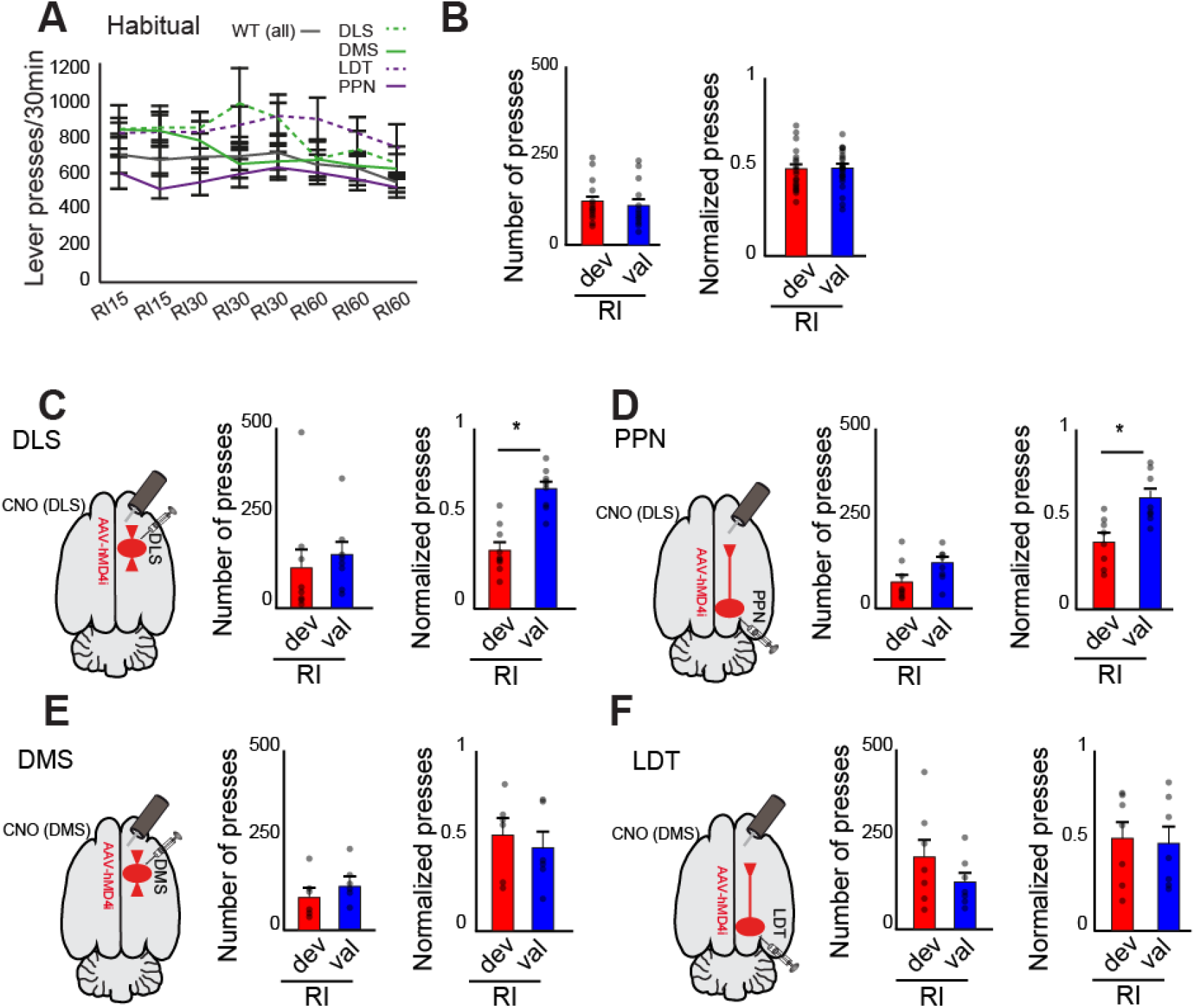
Blocking of cholinergic transmission in the dorsal striatum impairs habitual action control. A) Lever pressing during acquisition of habitual behavior shows no significant difference between groups (see text for details). (B-F) Number of presses and normalized lever presses during outcome devaluation testing across valued (val) or devalued (dev) states in random interval schedule (Rl; habitual). Control animals (WT, B) and ChAT::Cre animals were injected in the DLS (C), PPN (D), DMS (E) or LDT (F). During Rl, significant devaluation was observed In DLS and PPN groups (D and F), suggesting that inhibition of PPM and DLS CINs axons prevent animals from switching to habitual learning. Individual data points and mean ± SEM are shown. * *P* < 0.05.

Following RR training, WT animals showed a higher number of lever presses in the valued session compared to the devalued session with a significant effect on the session factor (two-way ANOVA group _[DLS, DMS, LDT, PPN]_ x session _[valued, devalued]_; F_group_(3,39) = 2.71, *P* = 0.0615; F_session_(1,39) = 365.33, *P* = 0.00001; F_interaction_ (3,39) = 0.73, *P* = 0.54). In contrast, following RI training, the same animals showed no significant differences in the number of lever presses in the valued and devalued sessions (two-way ANOVA: F_group_ (3,39) = 1.18, *P* = 0.333; F_session_(1,39) = 0.10, *P* = 0.7546; F_interaction_(3,39) = 0.26, *P* = 0.8534). To illustrate differences in the proportion of responses during the devaluation tests within subjects, we analyzed the normalized number of presses between test sessions (see Methods; **Fig. 5B, 6B**), and calculated the difference between valued and devalued responses as an index that reveals the ability of animals to adjust their responses after training in each schedule (**Fig. S11**). Following RR training sessions, control animals showed a strong preference to seek the instrumental outcome in the valued condition compared to the devalued condition, denoting a bias towards goal-directed behavior (two-way ANOVA group x condition: F_group_ (3,39) = 0.001, *P* = 1; F_session_(1,39) = 401.35, *P* = 0.00001, F_interaction_(3,39) = 0.82, *P* = 0.4927), whereas following RI training sessions, they did not show any preference, denoting a bias towards habitual behavior, thus revealing two fundamentally distinct forms of encoding action-outcome associations (two-way ANOVA group x session: F_group_ (3,39) = 1.73, *P* = 0.1803; F_session_(1,39) = 15.79, *P* = 0.0004, F_interaction_(3,39) = 2.23, *P* = 0.1036).

Next, we interrogated the contribution of acetylcholine to this behavior in two striatal regions (dorsolateral and dorsomedial) in the ChAT::cre animals compared to the WT animals. Because there was no difference in the number of lever presses between control groups (i.e. associated to the brain region targeted), the WT data was pooled into one single control group for the following analyses. We inhibited cholinergic transmission from axons arising in dorsolateral CINs (**Fig. 5C**), PPN (**Fig. 5D**), dorsomedial CINs (**Fig. 5E**) or LDT (**Fig. 5F**). No effect of group or interaction was observed in the response rates during RR training (two-way ANOVA group_[WT, DLS, DMS, LDT, PPN]_ x day: F_group_(4,199) = 2.21, *P* = 0.0696; F_day_(3,199) = 21.73, *P* = 0.00001; F_interaction_(12,199) = 0.35, *P* = 0.9780). After RR training, most animals showed a significant reduction in lever pressing in the devalued session relative to the valued session, similar to the reduction in WT (two-way ANOVA group x session: F_group_(4,99) = 3.48, *P* = 0.0109, F_session_(1,99) = 54.58, *P* = 0.00001; F_interaction_(4,99) = 4.52, *P* = 0.023). *Post hoc* pairwise comparisons (Tukey’s) revealed a significant difference in lever presses (devalued *vs* valued) in WT (P < 0.0001), DLS (P < 0.0001), DMS (*P* = 0.003) but not in LDT (*P* = 1) or PPN (*P* = 0.518) groups. The normalized number of presses also revealed a significant interaction (two-way ANOVA group x session, F_group_(4,99) = 0.001, *P* =1; F_session_(1,99) = 429.47, *P* = 0.00001 and F_interaction_(4,99) = 23.29, *P* < 0.00001) with post *hoc* pairwise comparisons (Tukey’s) showing significant effects in the WT (P < 0.0001), DLS (P < 0.0001), DMS (P < 0.0001) and PPN (P < 0.0001) but not LDT (*P* = 1) groups (**Fig. 5B-F**). Thus, animals in the LDT group showed virtually the same proportion of lever presses during both valued and devalued sessions suggesting reduced expression of goal-directed behavior (**Fig. 5F**). In other words, when acetylcholine release in the DMS arising from LDT terminals was disrupted during RR training, rats failed to associate the outcome with the instrumental action that produced it and were therefore insensitive to reward devaluation. This effect was evident by the absence of shift in the devaluation index in LDT despite the different training conditions (**Fig. S11**).

Finally, following retraining in the absence of CNO administration, we tested the effects of cholinergic transmission on habitual learning in the same group of animals (**Fig. 6**). During RI training (**Fig. 6A**), no effect of group, day or interaction was observed in the response rates (two-way ANOVA group x day: F_group_(4,399) = 2.30, *P* = 0.0581, F_day_(7,399) = 0.98, *P* = 0.4437; F_interaction_(28,399) = 0.36, *P* = 0.9991). Animals in the dorsomedial striatum and LDT groups showed no significant differences in the number of lever presses during the valued and devalued sessions, as controls did, suggesting that habitual behavior encoding remained intact (**Fig. 6E, F**) (two-way ANOVA: groups x condition_[valued *vs* devalued]_, F_group_(4,99) = 1.34, *P* = 0.26, F_condition_(1,99) = 0.445, *P* = 0.5019;F_interaction_(4,99) = 0.71, *P* = 0.58). However, there was a significant interaction in the normalized number of presses (two-way ANOVA groups x condition:F_group_(4,99) = 1.34, *P* = 1; F_condition_ (1,99) = 10.06, P = 0.0021; F_interaction_(4,99) = 5.79, P = 0. 0003), and *post hoc* pairwise comparisons (Tukey’s) revealed a significant difference of normalized lever press (devalued *vs* valued) in DLS (*P* = 0.001; **Fig. 6C**) and PPN (*P* = 0.018; **Fig. 6D**), but not for WT (*P* = 1; **Fig. 6B**), DMS (*P* = 0.316; **Fig. 6E**) and LDT (*P* = 1; **Fig. 6F**). This suggests that rats in the dorsolateral striatum and PPN groups failed to shift to a habitual responding state and remained goal-directed, as suggested by reduced lever-pressing in the devalued session compared to the valued session. Thus, reduced cholinergic transmission in the dorsolateral striatum, regardless of its origin (i.e., CINs and PPN), impairs the ability of rats to form habitual behavior, thus revealing that cholinergic neurons from the midbrain have a critical role in normal striatal operations.

## Discussion

Cholinergic transmission in the striatum powerfully modulates striatal output^46^, the activity of striatal interneurons^30,36,47^, the release of glutamate from cortical terminals^1^ and the release of dopamine from mesostriatal terminals^48,49^. We present here, detailed evidence of the functionality of a hitherto uncharacterized source of acetylcholine in the striatum originating in the midbrain. We show that cholinergic neurons of the PPN and LDT provide direct innervation of CINs and direct and indirect pathway neurons. We show that PPN and LDT axon terminals inhibit the activity of SPNs while activating CINs, suggesting a circuit mechanism in which PPN/LDT can modulate striatal activity through CINs. Finally, we show that inhibition of cholinergic transmission from either PPN, LDT or CINs impairs shifts in action control, suggesting that cholinergic transmission from the midbrain is necessary for normal encoding of behavior.

### Two functionally distinct cholinergic systems interacting in the striatum

Our results show that PPN and LDT cholinergic neurons inhibit the firing of SPNs with a similar magnitude as CINs, suggesting overlapping actions. Because PPN and LDT preferentially contact CINs over SPNs (as supported by both electron microscopy and monosynaptic rabies labeling), and because their latency for activating CINs is shorter than for inhibiting SPNs, we hypothesized that PPN/LDT neurons may be exerting their effects in the striatum partly through their connections with CINs. The inhibitory effects of the PPN/LDT over SPNs were significantly reduced but not abolished following the inhibition of CINs and completely abolished following the infusion of acetylcholine blockers. These results suggest that CINs play a key role in the modulation of the striatal output by PPN/LDT, but additional mechanisms are likely to contribute to the cholinergic modulation arising in the midbrain. Such mechanisms may rely on a monosynaptic modulation of SPNs by PPN/LDT cholinergic axons (as shown by our anatomical data in **Fig. 1** and the evidence of synaptic contacts with dendritic spines in^10^) or through other GABAergic interneurons (e.g.^31^).

The seemingly overlapping effects of cholinergic signaling arising from two different sources raise the question of whether they are conveying different messages or acting in coordination. Several differences between striatal CINs and midbrain cholinergic neurons have been reported in the literature. CINs receive innervation predominantly from cortical areas, including cingulate, secondary motor and primary somatosensory cortices^50^, and thalamic nuclei including the parafascicular and centrolateral^51,52^. Inputs to cholinergic midbrain neurons have not been fully identified, but largely differ from CINs as they predominantly arise in basal ganglia structures, including the substantia nigra pars reticulata and internal globus pallidus^53,54^; for a review see ^55^. In terms of the physiological properties, CINs possess a high-input resistance (200 MΩ) (for review see^4^) and have been associated with a spontaneous, tonically-active firing mode (3-10 Hz;^56^) that is mediated by inward rectifying potassium currents and a depolarization sag that induces rebound spike firing^57^. In contrast, PPN cholinergic neurons show a low firing rate *in vitro* (2-3Hz), a very high input resistance (600MΩ), display an A-current^58^, their firing seems to be modulated by M-currents^59^ and show fast-adaptive firing^18^. *In vivo,* identified PPN cholinergic neurons have been shown to fire phasically^18,19^. This evidence thus indicates that midbrain and striatal cholinergic cell groups differ in their afferent connectivity and physiological properties, suggesting they are modulated differently by their afferents and that their dynamics are distinct.

Another significant difference stems from electrophysiological recordings of putative striatal and midbrain cholinergic neurons in awake, behaving animals. Tonically-active neurons in the striatum encode a pause in their firing rate that is associated with behaviorally-relevant salient events^60,61^, which is correlated with the phasic activation of dopamine neurons in mesostriatal systems. Importantly, this pause is often preceded by a phasic increase in firing before the inhibition, mediated in part by thalamostriatal activation^35^ and followed by a rebound excitation. Neurons in the PPN, in contrast, increase their firing rate phasically during sensory cues that predict reward presentation^62^, presumably driving dopamine transients in the striatum. The multiphasic response of CINs during behaviorally-relevant salient events suggests the convergence of multiple synaptic drives that shape the burst-pause-rebound dynamics of CINs. The direct connectivity and excitatory nature of the midbrain input onto CINs suggest that PPN/LDT cholinergic neurons contribute to sculpting the response of CINs during behavior. Further experiments are needed to determine the extent of this modulation.

### Role of the PPN in adaptive behavior

Our data here also suggest that there are intersecting roles of CINs and PPN/LDT neurons in cholinergic-mediated striatal behavior. In the dorsolateral striatum, we revealed that inhibition of cholinergic signaling arising from either CINs or PPN neurons is able to block the transition from goal-directed to habitual behavior, whereas in the dorsomedial striatum inhibition of cholinergic signaling from LDT neurons is able to block goal-directed behavior. Together with our anatomical data showing preferential innervation of PPN and LDT neurons over CINs, these behavioral effects suggest that midbrain cholinergic neurons modulate the activity of CINs during behavioral switching and action control. Similar changes in the outcome of these tasks have been obtained following the interruption of the thalamostriatal projections that target CINs^40^ and corticostriatal projections^39^, or following excitotoxic striatal lesions^63^. All the above suggest that optimal encoding of behavioral information in the striatum is mediated by a series of factors that converge at the level of the CINs; furthermore, it reveals the role of the PPN as a key modulator of striatal activity through CINs.

In line with our findings, the role of the PPN in adaptive behavior and action control has been previously addressed by a series of experiments using lesions or pharmacological manipulations. For example, non-specific PPN lesions impair adaptation to incremental walking speeds in a motor task^64^, affect assimilation of new strategies with a consequent increase in perseverant responses^65,66^, and decrease the sensitivity to reward omissions^67^, thus denoting a failure in adjusting the behavioral state. Furthermore, pharmacological inhibition of the PPN produces a decrease in the responsiveness to degradation in contingencies between action and outcome, but did not change it if contingencies remain unchanged^68^, in line with findings showing impaired ability of rats to adapt to new strategies when the contingencies changed following inhibition of cholinergic transmission in the striatum^6,69^ or CINs lesions^70^. This body of evidence suggests that interrupting PPN activity has similar effects to those observed following disruption of cholinergic transmission in the striatum and raises the possibility that PPN is mediating such response. Our experiments here link these systems together by showing that interfering with cholinergic transmission in the striatum, regardless of its origin, has similar functional consequences for action control.

### Striatum as a main hub of PPN cholinergic projections

PPN and LDT have divergent ascending cholinergic projections that converge in the striatum following three different pathways. First, axon collaterals of cholinergic neurons innervate dopamine neurons in the SNc and VTA^71–
73^. Activation of this pathway leads to increased activity of dopamine neurons that project to the striatal complex^22^. Second, cholinergic neurons innervate thalamic nuclei that in turn project to the striatum. In particular, PPN densely innervates the parafascicular nucleus^74–76^, which in turn preferentially targets and modulates CINs^35,77^. Third, our results here reveal that PPN and LDT cholinergic neurons directly innervate both SPNs and CINs, with preferential innervation of the latter. The convergence of three different afferent systems arising from a single cell group in the midbrain puts the PPN in a key position as modulator of striatal activity and suggests that striatum (whether directly or indirectly) is the main target of cholinergic PPN projections, as no other PPN target receives such level of converging afferents from PPN cholinergic neurons. Furthermore, at least a proportion of these projections originate from the same neurons^10^, potentially indicating the simultaneous activation of dopamine, thalamic and striatal targets, and suggesting that these converging effects at the striatum level are inextricably linked.

What is the PPN signaling in the striatum and why is it relevant for behavior? PPN neurons have been shown to have a phasic activation during particular behavioral contexts, such as during Pavlovian conditioning^78^, reward prediction^79–80^ and reward omission^81^. Thus, when these signals are absent because PPN neurons fail to signal a mismatch between expected and real contingencies, the behavior is not updated, creating perseverant responses and failure to integrate new learning with the old learning (see^24^). The activation of CINs may thus underlie the mechanism by which PPN is able to shape striatal output and block ongoing motor programs at the level of SPNs in order to update the behavioral state and reinforce novel actions.

### Author contributions

Conceptualization, D.D. and J.M.S.; Methodology, D.D., I.H.O., M.V., K.K., T.G. and J.M.S.; Behavioral experiments developed in Leicester; Formal analysis, D.D., M.V. and J.M.S., Investigation, D.D. and I.H.O.; Writing – Original Draft, D.D. and J.M.S.; Writing – Review & Editing, D.D., I.H.O., M.V., K.K., T.G. and J.M.S.; Visualization, D.D., I.H.O., M.V., K.K., T.G. and J.M.S.; Supervision and Funding acquisition, J.M.S.

## Acknowledgments

We thank Paul Bolam for valuable input at different stages of this project. In addition, we also thank M. Shifflet and N. Gut for comments on this manuscript, M. Condon for some recordings in the initial stages of this project, and A.M. Aman for assistance in animal training. This research was supported by NIH grant R01 NS100824 (J.M.S.), a NARSAD Young Investigator Award (J.M.S.) and Rutgers University. MV acknowledges support from the Departamento de Salud, Gobierno de Navarra (114/2014) and Spanish Ministry of Education, Culture and Sport and Fulbright Commission (CAS15/00259).

## References

1. Malenka, R. C. & Kocsis, J. D. Presynaptic actions of carbachol and adenosine on corticostriatal synaptic transmission studied in vitro. J Neurosci 8, 3750–3756 (1988).

2. Pakhotin, P. & Bracci, E. Cholinergic interneurons control the excitatory input to the striatum. J Neurosci 27, 391–400 (2007).

3. Oldenburg, I. A. & Ding, J. B. Cholinergic modulation of synaptic integration and dendritic excitability in the striatum. Curr. Opin. Neurobiol. 21, 425–432 (2011).

4. Lim, S. A. O., Kang, U. J. & McGehee, D. S. Striatal cholinergic interneuron regulation and circuit effects. Front. Synaptic Neurosci. 6, (2014).

5. Aoki, S., Liu, A. W., Zucca, A., Zucca, S. & Wickens, J. R. Role of Striatal Cholinergic Interneurons in Set-Shifting in the Rat. J Neurosci 35, 9424–9431 (2015).

6. McCool, M. F., Patel, S., Talati, R. & Ragozzino, M. E. Differential involvement of M1-type and M4-type muscarinic cholinergic receptors in the dorsomedial striatum in task switching. Neurobiol Learn Mem 89, 114–124 (2008).

7. Graybiel, A. M., Baughman, R. W. & Eckenstein, F. Cholinergic neuropil of the striatum observes striosomal boundaries. Nature 323, 625–627 (1986).

8. Kubota, Y. & Kawaguchi, Y. Spatial distributions of chemically identified intrinsic neurons in relation to patch and matrix compartments of rat neostriatum. J. Comp. Neurol. 332, 499–513 (1993).

9. Matamales, M. et al. Quantitative Imaging of Cholinergic Interneurons Reveals a Distinctive Spatial Organization and a Functional Gradient across the Mouse Striatum. PLoS One 11, e0157682 (2016).

10. Dautan, D. et al. A major external source of cholinergic innervation of the striatum and nucleus accumbens originates in the brainstem. J Neurosci 34, 4509–4518 (2014).

11. Cornwall, J., Cooper, J. D. & Phillipson, O. T. Afferent and efferent connections of the laterodorsal tegmental nucleus in the rat. Brain Res Bull 25, 271–284 (1990).

12. Lerner, T. N. et al. Intact-Brain Analyses Reveal Distinct Information Carried by SNc Dopamine Subcircuits. Cell 162, 635–647 (2015).

13. Nakano, K. et al. Topographical projections from the thalamus, subthalamic nucleus and pedunculopontine tegmental nucleus to the striatum in the Japanese monkey, Macaca fuscata. Brain Res 537, 54–68 (1990).

14. Saper, C. B. & Loewy, A. D. Projections of the pedunculopontine tegmental nucleus in the rat: evidence for additional extrapyramidal circuitry. Brain Res. 252, 367–372 (1982).

15. Smith, Y. & Parent, A. Differential connections of caudate nucleus and putamen in the squirrel monkey (Saimiri sciureus). Neuroscience 18, 347–371 (1986).

16. Wall, N., DeLaParra, M., Callaway, E. & Kreitzer, A. Differential innervation of direct- and indirect-pathway striatal projection neurons. Neuron 79, 347–360 (2013).

17. Hunnicutt, B. J. et al. A comprehensive excitatory input map of the striatum reveals novel functional organization. Elife 5, (2016).

18. Petzold, A., Valencia, M., Pal, B. & Mena-Segovia, J. Decoding brain state transitions in the pedunculopontine nucleus: cooperative phasic and tonic mechanisms. Front Neural Circuits 9, 68 (2015).

19. Boucetta, S., Cissé, Y., Mainville, L., Morales, M. & Jones, B. E. Discharge profiles across the sleep-waking cycle of identified cholinergic, GABAergic, and glutamatergic neurons in the pontomesencephalic tegmentum of the rat. J. Neurosci. 34, 4708–27 (2014).

20. Cox, J., Pinto, L. & Dan, Y. Calcium imaging of sleep-wake related neuronal activity in the dorsal pons. Nat. Commun. 7, 10763 (2016).

21. Furman, M. et al. Optogenetic stimulation of cholinergic brainstem neurons during focal limbic seizures: Effects on cortical physiology. Epilepsia 56, e198–202 (2015).

22. Dautan, D. et al. Segregated cholinergic transmission modulates dopamine neurons integrated in distinct functional circuits. Nat. Neurosci. 19, 1025–1033 (2016).

23. Xiao, C. et al. Cholinergic Mesopontine Signals Govern Locomotion and Reward through Dissociable Midbrain Pathways. Neuron 90, 333–347 (2016).

24. Mena-Segovia, J. & Bolam, J. P. Rethinking the Pedunculopontine Nucleus: From Cellular Organization to Function. Neuron 94, 7–18 (2017).

25. Callaway, E. M. & Luo, L. Monosynaptic Circuit Tracing with Glycoprotein-Deleted Rabies Viruses. J. Neurosci. 35, 8979–8985 (2015).

26. Tepper, J. M., Abercrombie, E. D. & Bolam, J. P. Basal ganglia macrocircuits. Progress in Brain Research 160, 3–7 (2007).

27. Sharott, A., Doig, N. M., Mallet, N. & Magill, P. J. Relationships between the Firing of Identified Striatal Interneurons and Spontaneous and Driven Cortical Activities In Vivo. J.Neurosci. 32, 13221–13236 (2012).

28. Bertran-Gonzalez, J., Chieng, B. C., Laurent, V., Valjent, E. & Balleine, B. W. Striatal cholinergic interneurons display activity-related phosphorylation of ribosomal protein S6. PLoS One 7, e53195 (2012).

29. Lee, K. et al. Parvalbumin Interneurons Modulate Striatal Output and Enhance Performance during Associative Learning. Neuron 93, 1451–1463.e4 (2017).

30. English, D. F. et al. GABAergic circuits mediate the reinforcement-related signals of striatal cholinergic interneurons. Nat Neurosci 15, 123–130 (2012).

31. Faust, T. W., Assous, M., Tepper, J. M. & Koós, T. Neostriatal GABAergic Interneurons Mediate Cholinergic Inhibition of Spiny Projection Neurons. J. Neurosci. 36, 9505–11 (2016).

32. Roseberry, T. K. et al. Cell-Type-Specific Control of Brainstem Locomotor Circuits by Basal Ganglia. Cell 164, 526–537 (2016).

33. Xiao, C. et al. Chronic Nicotine Selectively Enhances 4 2* Nicotinic Acetylcholine Receptors in the Nigrostriatal Dopamine Pathway. J. Neurosci. 29, 12428–12439 (2009).

34. Ding, J., Peterson, J. D. & Surmeier, D. J. Corticostriatal and Thalamostriatal Synapses Have Distinctive Properties. J. Neurosci. 28, 6483–6492 (2008).

35. Ding, J. B., Guzman, J. N., Peterson, J. D., Goldberg, J. A. & Surmeier, D. J. Thalamic gating of corticostriatal signaling by cholinergic interneurons. Neuron 67, 294–307 (2010).

36. Faust, T. W., Assous, M., Shah, F., Tepper, J. M. & Koos, T. Novel fast adapting interneurons mediate cholinergic-induced fast GABAA inhibitory postsynaptic currents in striatal spiny neurons. Eur J Neurosci 42, 1764–1774 (2015).

37. Stachniak, T. J., Trudel, E. & Bourque, C. W. Cell-Specific Retrograde Signals Mediate Antiparallel Effects of Angiotensin II on Osmoreceptor Afferents to Vasopressin and Oxytocin Neurons. Cell Rep. 8, 355–362 (2014).

38. Gremel, C. M. & Costa, R. M. Orbitofrontal and striatal circuits dynamically encode the shift between goal-directed and habitual actions. Nat. Commun. 4, 2264 (2013).

39. Gremel, C. M. et al. Endocannabinoid Modulation of Orbitostriatal Circuits Gates Habit Formation. Neuron 90, 1312–1324 (2016).

40. Bradfield, L. A., Bertran-Gonzalez, J., Chieng, B. & Balleine, B. W. The thalamostriatal pathway and cholinergic control of goal-directed action: interlacing new with existing learning in the striatum. Neuron 79, 153–166 (2013).

41. Bradfield, L. A. & Balleine, B. W. Thalamic control of dorsomedial striatum regulates internal state to guide goal-directed action selection. J. Neurosci. 3860–16 (2017). doi:10.1523/JNEUROSCI.3860-16.2017

42. MacLaren, D. A. A., Wilson, D. I. G. & Winn, P. Selective lesions of the cholinergic neurons within the posterior pedunculopontine do not alter operant learning or nicotine sensitization. Brain Struct. Funct. 221, 1481–1497 (2016).

43. Halbout, B., Liu, A. T. & Ostlund, S. B. A closer look at the effects of repeated cocaine exposure on adaptive decision-making under conditions that promote goal-directed control. Front. Psychiatry 7, (2016).

44. Hilário, M. R. F. Endocannabinoid signaling is critical for habit formation. Front. Integr. Neurosci. 1, (2007).

45. Dickinson, A. Actions and Habits: The Development of Behavioural Autonomy. Philos. Trans. R. Soc. B Biol. Sci. 308, 67–78 (1985).

46. Yamamoto, K., Ebihara, K., Koshikawa, N. & Kobayashi, M. Reciprocal regulation of inhibitory synaptic transmission by nicotinic and muscarinic receptors in rat nucleus accumbens shell. J. Physiol. 591, 5745–5763 (2013).

47. Sullivan, M. A., Chen, H. & Morikawa, H. Recurrent inhibitory network among striatal cholinergic interneurons. J. Neurosci. 28, 8682–90 (2008).

48. Threlfell, S. et al. Striatal dopamine release is triggered by synchronized activity in cholinergic interneurons. Neuron 75, 58–64 (2012).

49. Cachope, R. et al. Selective activation of cholinergic interneurons enhances accumbal phasic dopamine release: setting the tone for reward processing. Cell Rep 2, 33–41 (2012).

50. Guo, Q. et al. Whole-Brain Mapping of Inputs to Projection Neurons and Cholinergic Interneurons in the Dorsal Striatum. PLoS One 10, e0123381 (2015).

51. Doig, N. M., Magill, P. J., Apicella, P., Bolam, J. P. & Sharott, A. Cortical and thalamic excitation mediate the multiphasic responses of striatal cholinergic interneurons to motivationally salient stimuli. J. Neurosci. 34, 3101–17 (2014).

52. Smith, Y. et al. The thalamostriatal system in normal and diseased states. Front. Syst. Neurosci. 8, 5 (2014).

53. Shink, E., Sidibe, M. & Smith, Y. Efferent connections of the internal globus pallidus in the squirrel monkey: II. Topography and synaptic organization of pallidal efferents to the pedunculopontine nucleus. J. Comp. Neurol. 382, 348–363 (1997).

54. Noda, T. & Oka, H. Distribution and morphology of tegmental neurons receiving nigral inhibitory inputs in the cat: an intracellular HRP study. J. Comp. Neurol. 244, 254–266 (1986).

55. Martinez-Gonzalez, C., Bolam, J. P. & Mena-Segovia, J. Topographical organization of the pedunculopontine nucleus. Front Neuroanat 5, 22 (2011).

56. Wilson, C. J., Chang, H. T. & Kitai, S. T. Firing patterns and synaptic potentials of identified giant aspiny interneurons in the rat neostriatum. J. Neurosci. 10, 508–19 (1990).

57. Goldberg, J. A. & Reynolds, J. N. Spontaneous firing and evoked pauses in the tonically active cholinergic interneurons of the striatum. Neuroscience 198, 27–43 (2011).

58. Takakusaki, K. & Kitai, S. T. Ionic mechanisms involved in the spontaneous firing of tegmental pedunculopontine nucleus neurons of the rat. Neuroscience 78, 771–794 (1997).

59. Bordas, C., Kovacs, A. & Pal, B. The M-current contributes to high threshold membrane potential oscillations in a cell type-specific way in the pedunculopontine nucleus of mice. Front Cell Neurosci 9, 121 (2015).

60. Apicella, P. The role of the intrinsic cholinergic system of the striatum: What have we learned from TAN recordings in behaving animals? Neuroscience 360, 81–94 (2017).

61. Apicella, P., Ravel, S., Deffains, M. & Legallet, E. The Role of Striatal Tonically Active Neurons in Reward Prediction Error Signaling during Instrumental Task Performance. J. Neurosci. 31, 1507–1515 (2011).

62. Okada, K., Toyama, K., Inoue, Y., Isa, T. & Kobayashi, Y. Different pedunculopontine tegmental neurons signal predicted and actual task rewards. J Neurosci 29, 4858–4870 (2009).

63. Hilario, M., Holloway, T., Jin, X. & Costa, R. M. Different dorsal striatum circuits mediate action discrimination and action generalization. Eur. J. Neurosci. 35, 1105–14 (2012).

64. MacLaren, D. A., Santini, J. A., Russell, A. L., Markovic, T. & Clark, S. D. Deficits in motor performance after pedunculopontine lesions in rats--impairment depends on demands of task. Eur J Neurosci 40, 3224–3236 (2014).

65. Wilson, D. I. G., MacLaren, D. A. A. & Winn, P. Bar pressing for food: differential consequences of lesions to the anterior versus posterior pedunculopontine. Eur. J. Neurosci. 30, 504–513 (2009).

66. Syed, A., Baker, P. M. & Ragozzino, M. E. Pedunculopontine tegmental nucleus lesions impair probabilistic reversal learning by reducing sensitivity to positive reward feedback. Neurobiol. Learn. Mem. 131, 1–8 (2016).

67. Leblond, M., Sukharnikova, T., Yu, C., Rossi, M. A. & Yin, H. H. The role of pedunculopontine nucleus in choice behavior under risk. Eur. J. Neurosci. 39, 1664–1670 (2014).

68. MacLaren, D. A. A., Wilson, D. I. G. & Winn, P. Updating of action-outcome associations is prevented by inactivation of the posterior pedunculopontine tegmental nucleus. Neurobiol. Learn. Mem. 102, 28–33 (2013).

69. Ragozzino, M. E., Mohler, E. G., Prior, M., Palencia, C. A. & Rozman, S. Acetylcholine activity in selective striatal regions supports behavioral flexibility. Neurobiol Learn Mem 91, 13–22 (2009).

70. Matamales, M. et al. Aging-Related Dysfunction of Striatal Cholinergic Interneurons Produces Conflict in Action Selection. Neuron 90, 362–373 (2016).

71. Bolam, J. P., Francis, C. M. & Henderson, Z. Cholinergic input to dopaminergic neurons in the substantia nigra: a double immunocytochemical study. Neuroscience 41, 483–494 (1991).

72. Gould, E., Woolf, N. J. & Butcher, L. L. Cholinergic Projections To the Substantia Nigra From the Pedunculopontine and Nuclei. 28, 611–623 (1989).

73. Oakman, S. A., Faris, P. L., Kerr, P. E., Cozzari, C. & Hartman, B. K. Distribution of pontomesencephalic cholinergic neurons projecting to substantia nigra differs significantly from those projecting to ventral tegmental area. J Neurosci 15, 5859–5869 (1995).

74. Sugimoto, T. et al. Cholinergic neurons in the nucleus tegmenti pedunculopontinus pars compacta and the caudoputamen of the rat: a light and electron microscopic immunohistochemical study using a monoclonal antibody to choline acetyltransferase. Neurosci Lett 51, 113–7. (1984).

75. Steriade, M., Pare, D., Parent, A. & Smith, Y. Projections of cholinergic and non-cholinergic neurons of the brainstem core to relay and associational thalamic nuclei in the cat and macaque monkey. Neuroscience 25, 47–67 (1988).

76. Pare, D., Smith, Y., Parent, A. & Steriade, M. Projections of brainstem core cholinergic and non-cholinergic neurons of cat to intralaminar and reticular thalamic nuclei. Neuroscience 25, 69–86 (1988).

77. Lapper, S. R. & Bolam, J. P. Input from the frontal cortex and the parafascicular nucleus to cholinergic interneurons in the dorsal striatum of the rat. Neuroscience 51, 533–45 (1992).

78. Yau, H.-J. et al. Pontomesencephalic Tegmental Afferents to VTA Non-dopamine Neurons Are Necessary for Appetitive Pavlovian Learning. Cell Rep. 1–12 (2016). doi:10.1016/j.celrep.2016.08.007

79. Hong, S. & Hikosaka, O. Pedunculopontine tegmental nucleus neurons provide reward, sensorimotor, and alerting signals to midbrain dopamine neurons. Neuroscience 282C, 139–155 (2014).

80. Okada, K., Toyama, K., Inoue, Y., Isa, T. & Kobayashi, Y. Different pedunculopontine tegmental neurons signal predicted and actual task rewards. J. Neurosci. 29, 4858–4870 (2009).

81. Tian, J. et al. Distributed and Mixed Information in Monosynaptic Inputs to Dopamine Neurons. Neuron 91, 1374–1389 (2016).

